# *Lactobacillus curvatus* Strains Specifically Show High Levels of Tolerance to Freeze-Thaw Stress

**DOI:** 10.1101/2020.02.04.934851

**Authors:** Ryunosuke Ikai, Koichi Tanabe, Jun Shima

**Author notes:** **Corresponding author:** Jun Shima, Faculty of Agriculture, Ryukoku University, 1-5 Yokotani, Seta Oe-cho, Otsu, Shiga 520-2194, Japan.

## Abstract

Tolerance to freeze-thaw stress is an important characteristic in recent fermentation processes. To gain insight into the freeze-thaw tolerance of lactic acid bacteria (LAB), we performed screening experiments and observed that several *Lactobacillus curvatus* strains showed high freeze-tolerance even in the absence of cryoprotectants. These *Lb. curvatus* strains also showed high levels of freeze tolerance in a milk fermentation process. *Lactobacillus sakei* (closely related to *Lb. curvatus*) was not revealed to be a freeze-thaw tolerant strain. These data indicate that *Lb. curvatus* has specific mechanisms underlying its tolerance to freeze-thaw stress.

**Importance:** Our findings demonstrate that *Lb. curvatus* strains frequently show high levels of freeze-thaw tolerance in both culture and milk and that *Lb. curvatus* strains are suitable as a model species for investigations of the molecular mechanisms underlying freeze-thaw tolerance in LAB and for applications in fermentation industries.

---

The responses of various microorganisms to environmental stress have drawn attention in both fundamental (1) and applied sciences including fermentation and medical sciences. In fermentation science, tolerance to environmental stress is a useful characteristic of microorganisms used in fermentation such as lactic acid bacteria (LAB) and yeast (2, 3), because such tolerance can improve fermentation processes. Freeze-thaw stress has become important to the more recently developed fermentation processes, as part of the efforts to improve the processes of the production and preservation of fermented foods including LAB (4). Freeze-thaw stress is a complex stress that can consist of dehydration, physical stressors, low temperature, and/or oxidative stresses (3).

To the best of our knowledge, mechanisms underlying the tolerance of LAB to freeze-thaw stress have not been reported. We conducted the present study to gain insight into freeze-thaw stress tolerance. It is known that cryoprotectants (e.g., glycerol and glutamate) improve the survivability of LAB after freeze-thaw stress (5, 6), and we thus decided to screen tolerant LAB strains under the absence of cryoprotectants.

We used approx. 70 LAB strains as the screening resource. These strains were originally isolated from fermented foods and plants in our laboratory. As a primary screening, we assessed the growth ability of the LAB strains after repeated freeze-thaw treatments. It was necessary to performed freeze-thaw treatments twice, because significant differences among the tested strains were not detected when a single freeze-thaw treatment was administered. We used *Lactobacillus delbrueckii* subsp. *bulgaricus* isolated from commercial yogurt as a control strain, and it showed approx. 0.1 optical density at 600 nm (OD_600_) in our experimental conditions. Among all 70 strains tested, 11 strains showed an OD_600_ >0.3; these 11 strains were used for the secondary screening.

In the secondary screening, we estimated the colony forming units (CFUs) after a freeze-thaw treatment of the 11 stains selected in the primary screening (Fig. 1). The CFUs of each strain before and after the freeze-thaw treatment (Fig. 1A) revealed that before the freeze-thaw treatment, the CFUs of the strains ranged from 5 × 10^6^ to 5 × 10^8^. The CFU value of the control strain *Lb. delbrueckii* subsp. *bulgaricus* after the freeze-thaw treatment was greatly reduced (by a factor of 10^2^). Figure 2B illustrates the survivability ratio, i.e., the percentage of CFU after the freeze-thaw treatment to the CFU before treatment. The survivability ratios of four strains (IR1, IR7, IR8, and IR28) were >60%.

**Fig. 1.**
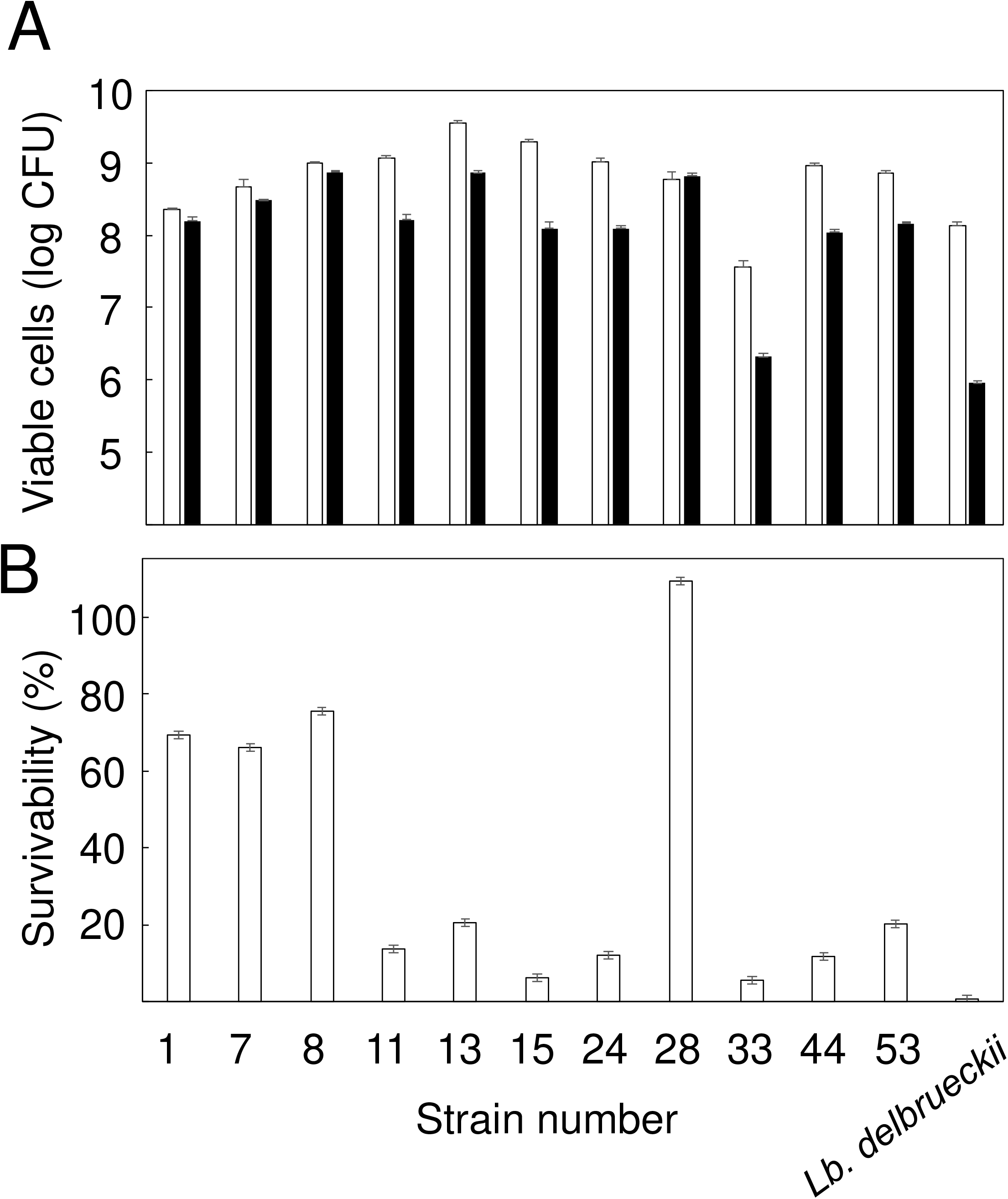
Survivability of LAB strains in culture. **A:** CFUs before and after freeze-thaw treatment. *White bars* and *black bars* present the CFUs before and after repeated freeze-thaw treatments, respectively. The data are average ± SD. **B:** Survivability is expressed as percent ± SD.

**Fig. 2.**
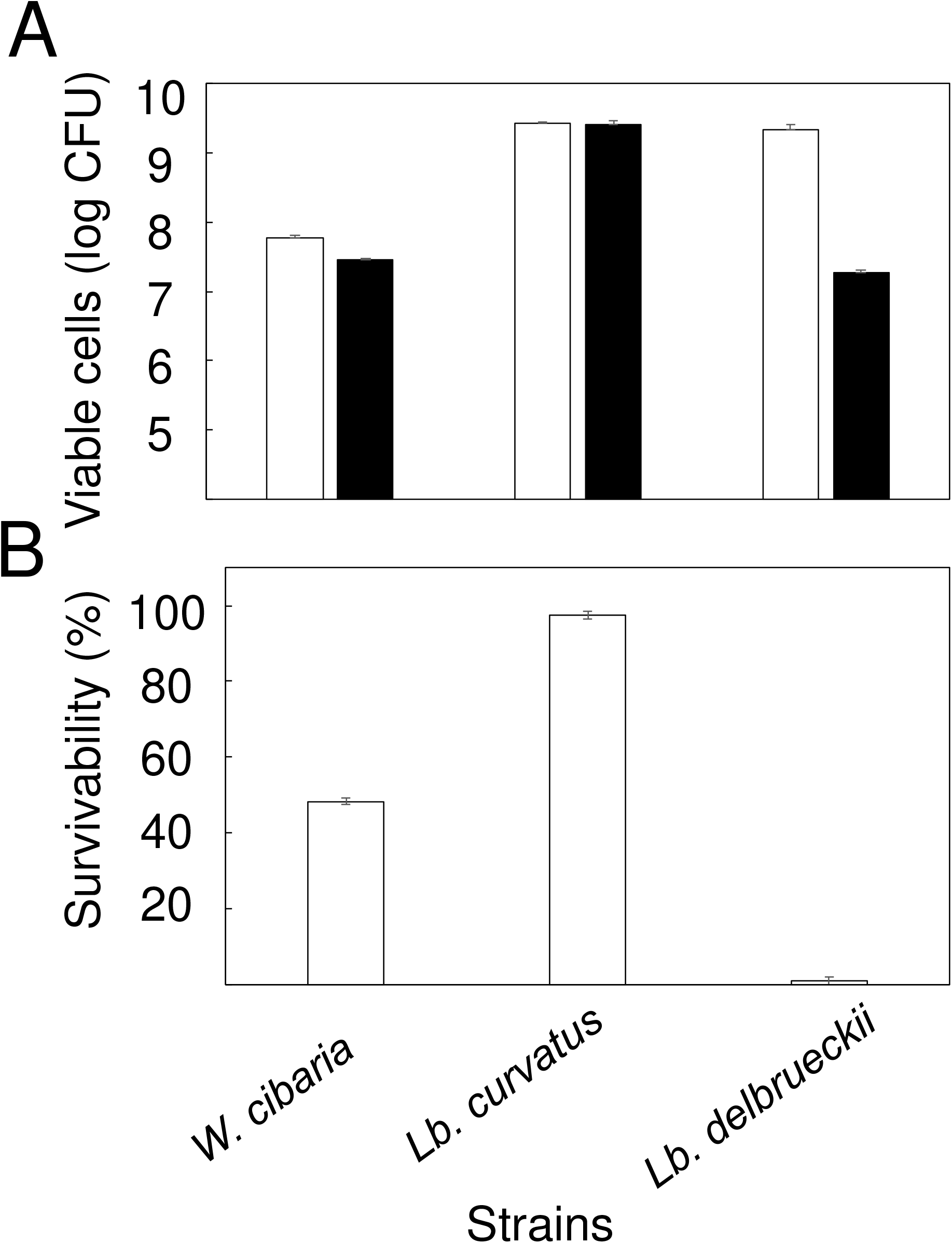
Survivability of LAB strains in milk. **A:** CFUs before and after freeze-thaw treatment. *White bars* and *black bars* present the CFUs before and after freeze-thaw treatments, respectively. The data are average ± SD. **B:** Survivability is expressed as percent ± SD.

We performed a taxonomic identification of the four strains based on the 16S rDNA sequence. Partial sequencing of 16S rDNA and the results of a BLAST analysis suggested that IR7, IR8, and IR28 are strains belonging to the species *Lb. curvatus* and that IR1 belongs to the species *Weissella cibaria* (Table S1).

We perform an analysis of the survivability of the *Lb. curvatus* IR8 and *W. cibaria* IR1 strains in milk that contained a potential cryoprotectant, because LAB strains are often used to produce fermented foods derived from milk products such as yogurt. After the addition of cells of *Lb. curvatus*, we observed strong coagulation of the milk (data not shown). In contrast, partial milk coagulation observed when *W. cibaria* IR1 was used under the same conditions. As shown in Figure 2, *Lb. curvatus* IR8 showed much higher survivability compared to *W. cibaria* IR1 and *Lb. delbrueckii* subsp. *bulgaricus*. These data suggested that *Lb. curvatus* strains may be useful in the production of fermented foods derived from milk.

*Lb. curvatus*, which are closely related to *Lactobacillus sakei* (7), is detected in a variety of resources including fermented foods and animal intestine, and *Lb. curvatus* has shown genomic diversity (8, 9). Our present analyses identified three *Lb. curvatus* strains with freeze-thaw tolerance. The three strains were isolated from different isolation resources Although *Lb. sakei* was included in our screening resources (data not shown), no *Lb. sakei* strain was detected as a freeze-thaw-tolerant strain.

The *Lb. curvatus* strains screened in this study may produce extracellular polysaccharide (EPS) (data not shown), and it is possible that EPS functions in cell protectants under freezing conditions (10). Although the molecular mechanisms underlying tolerance to freeze-thaw stress are not yet known, it is possible that *Lb. curvatus* has specific genes and/or cellular systems that contribute to the tolerance observed in three strains.

## Strains and medium

The LAB strains (~70 strains) had been preserved in our laboratory. They were obtained from natural resources in Japan including fermented foods. *Lb. delbrueckii* subsp. *bulgaricus* isolated from a commercial yogurt product (Meiji, Tokyo) was used as a control strain. All LAB strains were cultivated at 30°C in MRS medium (BD Biosciences, San Jose, CA) under anaerobic conditions provided by an AnaeroPack (Sugiyama-gen Co., Tokyo).

## Primary screening for freeze-tolerant LAB strains

The primary screening for freeze-tolerant LAB strains was performed based on the strains’ growth ability after freezing-thawing treatment. LAB strains were cultivated for 48 h in 5 ml of MRS broth. The cultures were frozen for 4 d at −25°C and thawed at 30°C, and they were then subjected to the same −25°C freeze for 4 days followed by thawing at 30°C. Portions (100 μL) of the samples were inoculated into 5 mL of fresh MRS broth and cultivated for 48 h. The OD600 was measured.

## Secondary screening for freeze-tolerant LAB strains

The secondary screening was performed to determine the strains’ viability, which was evaluated as CFUs. The LAB strains that were selected by the primary screening were cultivated for 48 h in 5 ml of MRS broth. The cultures were frozen for 4 d at −25°C and then thawed at 30°C. The measurement of the samples’ CFUs after the freezing and thawing treatment was conducted by first spreading the samples onto MRS agar medium in triplicate. After 48 h incubations, the numbers of viable colonies were counted. As control experiments, the CFUs were also measured in the samples before the freezing.

## Taxonomic identification of the screened strains

The taxonomic identification of the LAB strains was performed based on a partial sequence of 16S rDNA by the method described by Kawamoto et al. (11). We determined the homology of the sequences by conducting a BLAST search of the DNA Data Bank of Japan (DDBJ). The sequences of the LAB strains obtained in this study were deposited in the DDBJ. The accession numbers for the 16S rDNA sequences are as follows. *L. mesenteroides* IR1: LC520099. *Lb. curvatus* IR7: LC520100. *Lb. curvatus* IR8: LC520101. *Lb. curvatus* IR28: LC520102.

## The survivability of the screened strains after freezing and thawing in milk

To gain insights into potential commercial uses of the selected strains, we investigated the screened LAB strains’ survivability in milk. The LAB strains were cultivated for 48 h in 5 ml of MRS broth. The cells were collected by centrifugation (3,000 *g* for 10 min), washed with saline, and then added into 10 mL of milk containing 3.6% lipid (Meiji) and incubated for 24 h at 30°C. The fermented milk was frozen for 3 d at −25°C. The survivability of the cells in milk or solidified milk was estimated by the pour plate method. In brief, portions of the fermented milk were mixed with MRS agar medium and then poured into petri dishes in triplicate. After 48 h incubations, viable colonies were counted.

## ACKNOWLEDGEMENT

We declare that we have no conflicts of interest.

---

**Suppl. Table S1.** Summary of the BLAST search results of the screened LAB strains

| Strain | Homology (accession no. of reference sequence) |
| --- | --- |
| IR1 | 100% with *Weissella cibaria* LMG 17699<sup>T</sup>(NR_036924) |
| IR7 | 100 % with *Lactobacillus curvatus* JCM 1096<sup>T</sup> (LC063167) |
| IR8 | 100 % with *Lactobacillus curvatus* JCM 1096<sup>T</sup> (LC063167) |
| IR28 | 100 % with *Lactobacillus curvatus* JCM 1096<sup>T</sup> (LC063167) |

